# Interplay of Driver, Mini-Driver, and Deleterious Passenger Mutations on Cancer Progression

**DOI:** 10.1101/084392

**Authors:** Xin Li, D. Thirumalai

## Abstract

Cancer is caused by the accumulation of a critical number of somatic mutations (drivers) that offer fitness advantage to tumor cells. Moderately deleterious passengers, suppressing cancer progression, and mini-drivers, mildly beneficial to tumors, can profoundly alter the cancer evolutionary landscape. This observation prompted us to develop a stochastic evolutionary model intended to probe the interplay of drivers, mini-drivers and deleterious passengers in tumor growth over a broad range of fitness values and mutation rates. Below a (small) threshold number of drivers tumor growth exhibits a plateau (dormancy) with large burst occurring when a driver achieves fixation, reminiscent of intermittency in dissipative dynamical systems. The predictions of the model, in particular the relationship between the average number of passenger mutations versus drivers in a tumor, is in accord with clinical data on several cancers. When deleterious drivers are included, we predict a non-monotonic growth of tumors as the mutation rate is varied with shrinkage and even reversal occurring at very large mutation rates. This surprising finding explains the paradoxical observation that high chromosomal instability (CIN) correlates with improved prognosis in a number of cancers compared with intermediate CIN.

---

Cancer is an evolutionary disease caused by genomic instability resulting from sequential somatic mutations accumulated over the lifetime of an individual^1–3^. These mutations arise due to both exogenous (for example, DNA damage due to UV radiation and harmful environmental factors) and endogenous factors (such as errors during mitosis or epigenetic alterations). It is believed that virtually all manifestations of cancer are due to accumulation of such mutations in oncogenes and tumor suppressor genes^4^. The development of next-generation sequencing technologies has made whole-genome sequencing possible^5^. These *tour de force* studies have revealed the occurrence of hundreds or even tens of thousands of genetic mutations in a variety of cancers^6–10^. Most of the somatic mutations are thought to be benign, but steady accumulation of such mutations during the course of a lifetime could have deleterious consequences. Among the large number of somatic mutations, a few, involved in the activation of oncogenes^11,12^ and dysfunction of tumor suppressor genes^13,14^, play a critical role in driving cancer progression. Such mutations are the major “drivers”^5^. The rest of the genetic alterations are “passengers”, which are generally assumed to merely accompany the driver mutations without a significant role in tumor growth^15,16^.

The drivers bestow selective growth advantage to tumor cells. Because of their singular importance in driving tumor growth, the identification of these crucial mutations from a large number of somatic mutations has been one of the major goals in cancer research^17^. However, the problem is exacerbated because driver mutations in some of the oncogenes occur only infrequently. Along with the drivers, thousands of “passenger” mutations (passengers from now on) also accumulate spontaneously in the genome. Much less attention has been paid to the large population of passengers, which are usually assumed to be totally neutral or inconsequential on cancer progression. However, in two recent insightful studies^18,19^, it was found that the passengers cannot be ignored because they could have moderately deleterious effect on cancer cells. They found that the collective effect of deleterious passengers could overwhelm the effect of drivers even if individually they have negligible impact^18,19^. The rapid accumulation of deleterious passengers could even drive tumor cell population to extinction under certain conditions, although with enough drivers proliferation of cancer cells would eventually occur. In light of these interesting predictions, which apparently have experimental support^20^, we find it necessary to further investigate the role of other types of passengers in cancer progression.

Although regarded as neutral, moderately deleterious passengers could also get fixation, and hence influence the growth dynamics of cell populations^18,21^. In the same spirit, it has been proposed that mildly beneficial passengers (or “mini-drivers”) could also offer fitness advantages to tumors^22,23^. It is likely that the cell population would go extinct as weakly selected, moderately deleterious mutations accumulate continuously, which is the Muller's ratchet effect^24,25^. The presence of mildly beneficial mutations can counteract this effect because even a small fraction of mini-drivers is sufficient to maintain cellular homeostasis^26^.

Recently Castro-Giner et.al.^23^ have emphasized the important role mini-drivers might play in promoting tumor growth. Here, we quantitatively investigate the consequence of the interplay between driver, mini-driver, and deleterious passenger in the dynamics of tumor progression. Using an extension of the recently proposed model^18,19^, we analyze systematically distinct somatic mutations and their interactions in order to elucidate the mechanism of cancer progression. Our theoretical study, which belongs to a class of mathematical description of tumor growth based on evolutionary dynamics^27^, leads to the following predictions: (i) Mini-drivers, even if present in small numbers, can offset the effects of deleterious passengers and maintain population homeostasis as long as the number of drivers is small. (ii) If the number of drivers remains below a threshold value, the neutralizing effect of mini-drivers and deleterious passengers drive the tumor to dormancy for a period of time. (iii) Using distinct choices for the dependence of the cell death rate on the size of the population, we find that the number of drivers leading to unbounded growth is ~ 4, which is similar to a recent estimate^28^. (iv) The predicted relationship between the total number of passengers as a function of driver mutations is in very good agreement with clinical data on several cancers. Interestingly, this relationship varies greatly from one trajectory to another, providing insights into the origins of cancer heterogeneity. (v) Our work also shows that inclusion of deleterious drivers results in reversal of tumor growth at high mutation rates (> 10^−5^), which tidily explains the seemingly paradoxical correlation between high chromosomal instability and improved prognosis in a variety of cancers.

## RESULTS

In principle, five distinct genetic mutations could appear in daughter cells when cells divide. Drivers and passengers are mutations which exert strong and weak influences on cell proliferation, respectively. These mutations could be beneficial or deleterious, which separately promote or decrease cell reproduction. The four types of somatic mutations are beneficial drivers (Ds), deleterious drivers (DDs), beneficial passengers (or mini-drivers (MDs)), and deleterious passengers (DPs). The fifth is a neutral genetic alteration with negligible effect on cancer progression.

Although all five types of mutations could occur in somatic cells, their influences on cancer development vary greatly. Neutral passengers can be neglected because they have no influence on cell proliferation or cell death. The deleterious driver (DDs) either slows down the proliferation of cells or increases their death rate strongly, leading eventually to rapid extinction of cells. We first disregard DDs because they are unlikely to get fixation in the population but will discuss their potential role at the end of this article.

Because a large number of passengers have been discovered in cancer cells that accompany driver mutations, they are likely to play an important role during cancer progression even though their effects have not been systematically studied. Simulations based on the tug-of-war model^18,19^ show that DPs and Ds work against each other during cancer progression. The results suggest that passengers could even overwhelm the effects of drivers provided the initial tumor size is not too large, resulting in reversal or even elimination of tumor progression. This finding contradicts the standard lore, which assume that the influence of passengers could be totally ignored^29^. The relevance of MDs has been discussed in the context of asexual populations where it is found that such mutations are important in maintaining population homeostasis^26^. In the absence of such mutations cell population would go extinct rapidly because of the Muller’s ratchet effect^24,25^. However, the relevance of mini-drivers, whose importance has been emphasized only recently^23^, has not been quantitatively assessed in previous evolutionary models dealing with cancer progression.

We first consider the interplay of DPs, MDs, and Ds in controlling cancer progression, with primary focus being the effect of MDs on tumor growth. A single driver mutation confers a fitness advantage s_d_. Similarly, a cell acquires a fitness advantage *s*_*md*_ or disadvantage *s*_*dp*_ as a single mini-driver or deleterious passenger accumulates, respectively. The fitness values satisfy the inequality *s_d_* ≫ *s*_*md*_, *s*_*dp*_ because the impact of driver mutations is much stronger than the mini-drivers and passengers. For simplicity, we choose *s*_*md*_ = *s*_*dp*_ ≡ *s*_*p*_ for the fitness of MDs and DPs in most of the simulations. The consequences of relaxing this assumption is discussed in the Supplementary Information (SI). It may appear that the inclusion of mini-drivers should merely renormalize the fitness advantage of the drivers.

However, because the frequency of accumulation of Ds and MDs and their numbers are vastly different, this is not the case as we establish here.

Tumor progression is based on evolutionary dynamics described in Fig. 1. A cell divides at a rate *B(d, md, dp)* where *d, md* and *dp* are the number of Ds, MDs and DPs, respectively. The only requirement is that *B*(*d,md,dp*) should be an increasing function of d and *md*, and should decrease as *dp* increases. The following form *B*(*d, md, dp*), a generalization of the one used previously^18^,

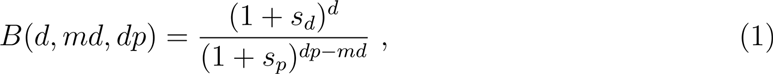

satisfies the criterion stated above. It is likely that the chosen functional form for *B*(*d, md, dp*) may not affect the overall growth kinetics but the details might differ depending on the precise form of *B*(*d, md, dp*). For comparison with previous work^18^, we assume that the death rate *D*(*N*) is given by,

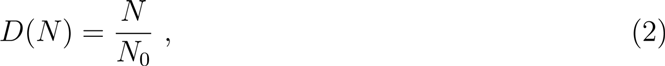

where *N*_0_ is the initial size of the population without mutations. Other forms of *D*(*N*) are also considered below. Cells acquire genetic mutations at a rate *μ* per locus after division. The numbers of driver, beneficial, and deleterious passenger loci are *L_d_*, *L*_*md*_, *L*_*dp*_, which satisfy *L*_*d*_ ≪ (*L*_*md*_ + *L*_*dp*_) ≡ *L*_*p*_. Thus, the frequency of driver and passenger mutations per cell division are given by *μL_d_*, *μL*_*md*_, and *μL*_*dp*_ as illustrated in Fig. 1, which also shows the kinetic scheme for cancer progression. Unless otherwise stated we use the numerical parameters in Table 1.

**FIG. 1.**
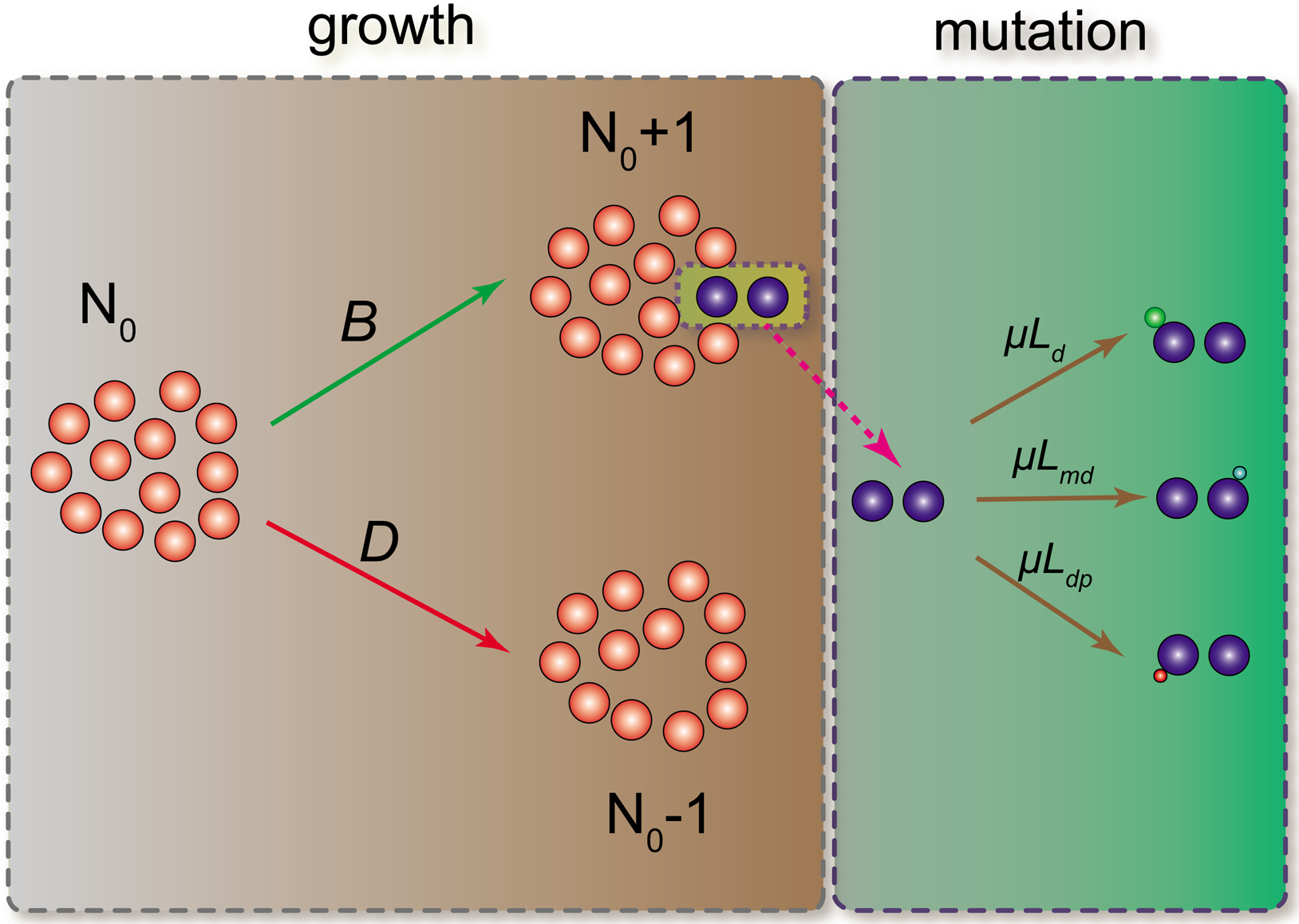
Growth and mutation of cancer cell starting from an initial population, *N*_0_. Each cell divides stochastically with birth rate *B(d,md,dp)* (Eq. (1)). A cell can also die stochastically with death rate *D*(*N*) (Eq. (2)). Cells accumulate driver, mini-driver and deleterious passenger mutations with rates *μL_d_*, *μL*_*md*_, and *μL_dv_* respectively, after each cell division. A single driver mutation increases the cell viability by *S_d_* while a mini-driver (deleterious passenger) confers fitness advantage (disadvantage) by *s_md_* (*s_dp_*).

**TABLE I.** Parameters for dynamics of cancer progression if not mentioned specifically. The value for *L_dd_* is estimated in this work and the values of other parameters are taken from previous studies^18^.

| parameter | value |
| --- | --- |
| $\mu$, mutation rate | $10^{-8}$ |
| $L_d$, number of driver loci | 1400 |
| $L_p$, number of passenger loci | $10^7$ |
| $L_{dd}$, number of deleterious driver loci | $10^4$ |
| $s_d$, fitness increase for one driver mutation | 0.1 |
| $s_p$, fitness decrease for one passenger mutation | 0.001 |

The stochastic growth for cancer progression is implemented using the Gillespie algorithm^30^ with defined reaction rates given in Fig. 1. We consider four distinct variations starting with a model, taking into account only the effect of DPs In the second case, we consider both DPs and MDs. The third, previously investigated variation^18^, examines the influence of both drivers and deleterious passengers. Finally, we investigate the new model probing the interplay of Ds, MDs, and DPs in driving cancer growth. A systematic investigation of the four cases allows us to isolate the influence of each of mutation type on cancer growth and adaptation.

**Interplay of deleterious passengers and mini-drivers lead to population homeostasis.** In contrast to the results in Fig. S1, the population dynamics changes dramatically when the effect of MDs are also taken into account. In a different context, it has been shown that beneficial passengers are required for the maintenance of a stable asexual population over a prolonged period of time^26^. Mini-drivers, which confer fitness advantage to cancer cells also accumulate during the progression of cancers as a result of stochastic genetic mutations^22,23^. The total number of loci (*L*_*p*_) for passenger mutations is fixed. A small fraction of such mutations is MDs. The number of MDs changes with time and can only be determined from the trajectories generated using the evolutionary dynamics described in Fig. 1. A generic evolutionary model^31^ probing the fate of finite population shows that the frequency of acquiring beneficial mutations is not a constant, but decreases as the fitness of cells increases. A corollary of this observation is that deleterious mutations accumulate more readily as the fitness of cells increases. From Eq. (1), we note that the fitness of cells is determined by the difference md – *dp* in the absence of drivers. We assume a linear relation for the fraction *F*_MD_, which is proportional to the probability of acquiring a MD upon cell division, yielding,

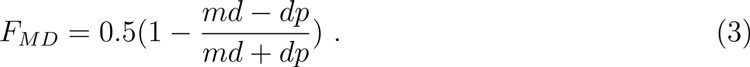

Note that *F*_*MD*_ decreases as cell fitness increases. In the presence of both DPs and MDs, Fig. 2A shows that *N*(*t*) ≈ *N*_0_ is time independent. The population is not driven to extinction but reaches homeostasis; *N*(*t*) fluctuates around a constant value for long periods of time (Fig. 2A).

**FIG. 2.**
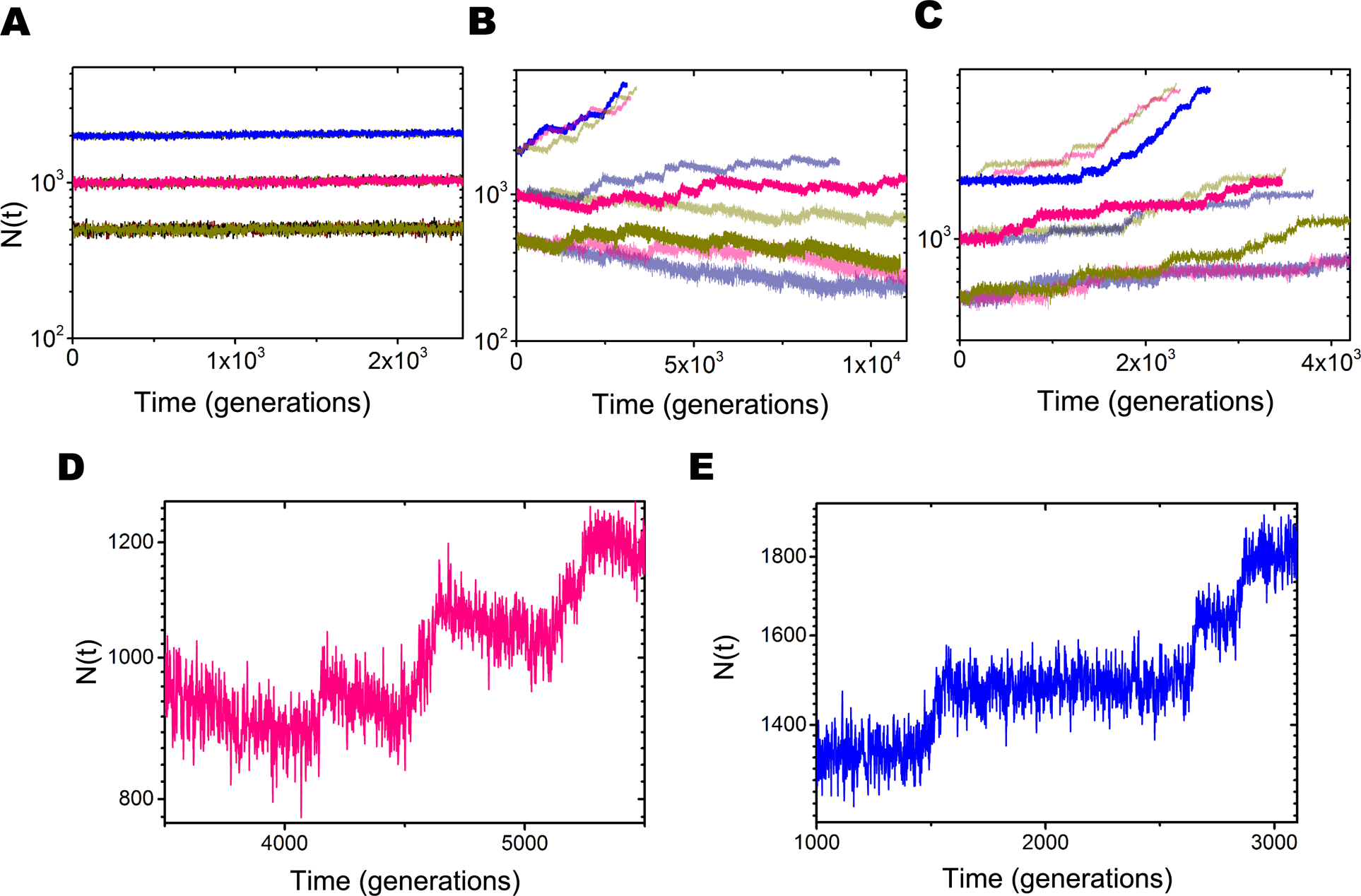
Population size as a function of time (generations) for different models. A generation is roughly three days. (A) Results for a model with deleterious and beneficial passengers. (B) Effect of deleterious passengers and drivers on tumor growth. (C) Interplay between drivers, deleterious passengers as well as mini-drivers on *N*(*t*). (D) A plot of *N*(*t*) over a short period for the red trajectory in (B). (E) Same as (D) except this plot is for *N*(*t*) in blue in (C). For each case three trajectories, with the initial population size *N*_0_ = 500, 1000, or 2000, are shown. The parameter values are taken from Table 1.

In order to assess the robustness of the finding that interplay of DPs and MDs confers homeostasis (Fig. 2A), we also performed simulations using another form of *F*_*MD*_ (see SI for details). An example displayed in the inset to Fig. S2 shows that a small fraction of MDs is sufficient to maintain homeostasis over a wide parameter range. The results in Fig. 2A and the inset of Fig. S2 show that homeostasis can be maintained although passenger mutations accumulate continually. This is accomplished through regulation of the ratio of MDs to DPs. However, it is believed that during cancer progression, homeostasis is intermittently halted leading to the population growth and eventual occurrence of the disease due to the accumulation of driver mutations.

**Drivers qualitatively alter growth of tumors.** The case without MDs has been investigated previously^18^ using a model in which cells accumulate only DPs and driver mutations. The growth trajectories are quite different even under the same initial conditions (Fig. 2B), due to the stochastic occurrence of driver mutations. Once a driver mutation sweeps through the whole population, *N*(*t*), increases rapidly. However, the system size shrinks gradually before the appearance of an additional driver mutation, which is seen in *N*(*t*) curves as an inverse sawtooth pattern (Fig. 2D). Thus, there is a tug-of-war between the beneficial driver mutations and DPs, with the former overwhelming the latter if there is sufficient number of drivers. Interestingly, there is a critical population size, *N*_*_, in this model with the unbounded growth occurring above *N*_*_ and extinction of *N*(*t*) below it (Fig. 2B). If the fixation frequency of driver mutations becomes very low, the whole population would be eliminated because of the accumulation of DPs. This occurs in the long time limit only for small values of *N*_0_, prompting a suggestion that a plausible route to therapy is to enhance the rate of DPs production. On the other hand, drivers would win and lead to the occurrence of cancer as they get fixation frequently when *N*_0_ exceeds *N*_*_.

**Intermittency in cancer growth is driven by mini-drivers.** Although the tug-of-war model revealed the importance of DPs, the very existence of *N*_*_ have not been documented in tumor evolution. In addition, *N*(*t*) does not go extinct if driver mutations appear rarely or not at all, indicating that both the driver mutations and MDs are crucial for explaining the dynamics of cancer progression. In our model that also includes MDs, we observe a totally different growth dynamics (Fig. 2C). For most of the time, homeostasis is maintained with very small variations in *N*(*t*) until the appearance of a single driver mutation. Once the driver mutation sweeps the whole population, *N*(*t*) increases rapidly (see Fig. S4 in the SI) until the death rate catches up with the higher fitness gained from a driver mutation. This leads to another period of homeostasis as illustrated by plateaus in Fig. 2E. The population does not shrink between fixation of two successive driver mutations. In contrast to the results in Fig. 2B, *N*(*t*) does not collapse if drivers appear rarely. However, continuous accumulation of driver mutations eventually disrupts homeostasis resulting in an unbounded growth (Fig. 2C) independent of *N*_0_. In other words, a threshold *N*_*_ does not exist in our model.

The evolution of *N*(*t*) resembles intermittency phenomenon in dissipative dynamical systems when a control parameter is altered. In our model such a parameter is the number (d) of drivers. When d is below a threshold value, *N*(*t*)exhibits an intermittent growth. When the threshold is exceeded there is an explosive growth in *N*(*t*)that is reminiscent of transition to chaos^32^.

Even with the inclusion of MDs, the initial population size is important for cancer progression, as illustrated in Fig. 2C. It takes much less time for the tumor to reach a macroscopic size if *N*_0_ is relatively large. The population does not go extinct if *N*_0_ is small, but it takes longer time for cancer to occur. The robustness of the results is established in Fig. S2 by considering an alternative functional form for *F_MD_*. Cancer dormancy is often observed during tumor progression^33,34^, and the homeostasis in Fig. 2C and Fig. S2 explains this observation, which in our model is a consequence of MDs. A dynamic equilibrium is maintained by the three distinct types of mutations during the period of dormancy until the occurrence of a new driver, which inevitably drives the tumor to a new stage of development. After a sufficient number of drivers accumulate there is an explosive increase in *N*(*t*).

**Tumor growth rate increases with mutation rate.** Typically, cancers appear more rapidly upon exposure to chemical carcinogens or when the mutation rate *μ* is increased either by environmental fluctuations or genetic mutations at late stages^3^. Interestingly, in the tug-of-war model^18^, it is found that the probability of cancer is low if *μ* exceeds a critical value, *μ_c_*. At the smallest *μ* (= 10^−9^) the increase in *N*(*t*)is slow (orange in Fig. 3A) whereas at *μ* = 10^−8^ the growth is more rapid as the driver mutations get fixation. For the parameter values listed in Table 1 and with *N*_0_ = 2000, the population grows as driver mutations accumulate (Fig. 2B and the red line in Fig. 3A). However, at a higher *μ*, *N*(*t*)shrinks as time progresses with *N*(*t*) → 0 (blue curve in Fig. 3A).

**FIG. 3.**
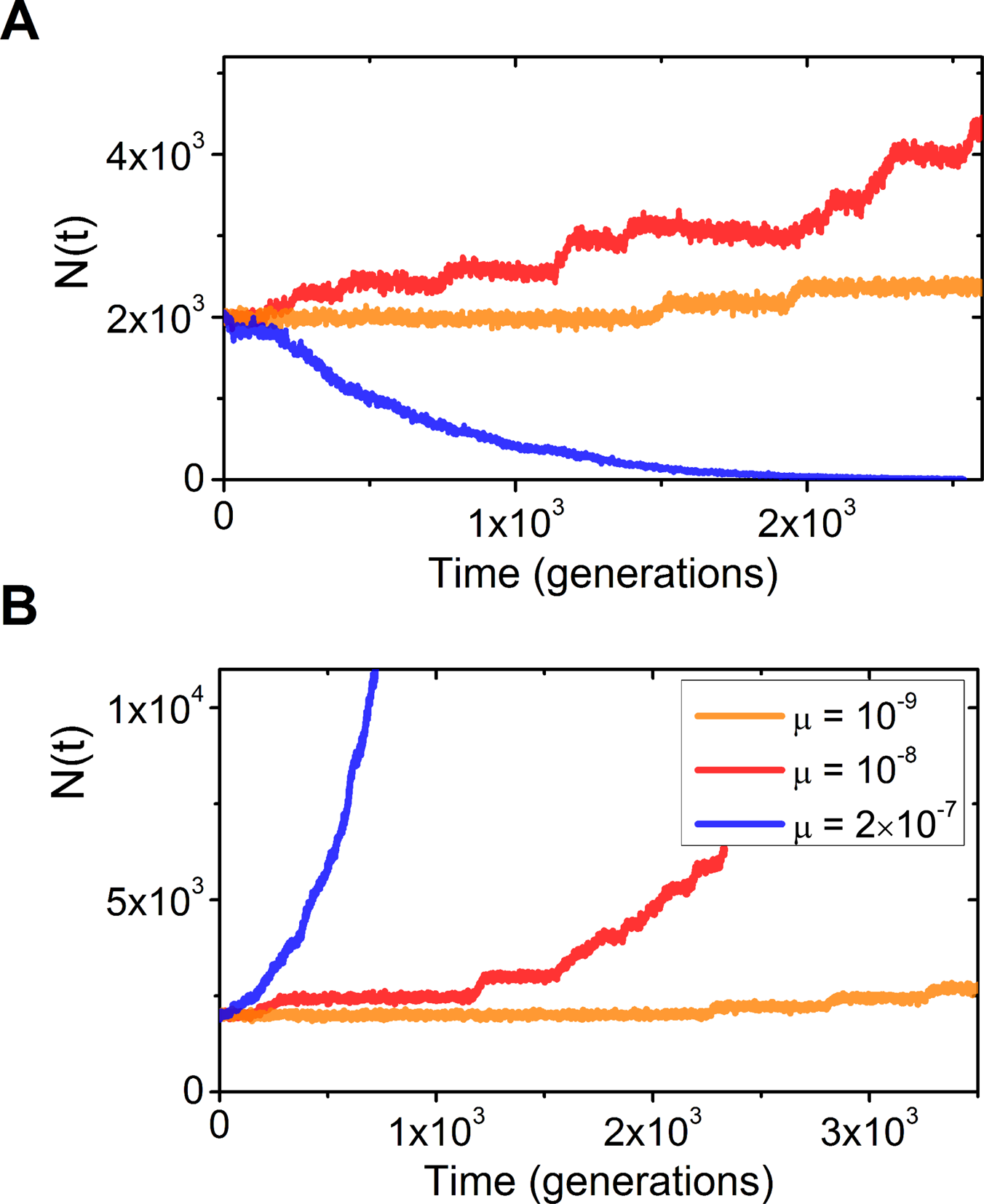
Population size as a function of time (generations) at three mutation rates *μ*. (A) Tug-of-war model with deleterious passengers and drivers. The *μ* values are shown in the inset of (B). (B) Model including deleterious, beneficial passengers as well as driver mutations. Besides *μ* all other parameter values are listed in Table 1.

Inclusion of MDs leads to qualitatively different results. There is complete absence of *μ_c_* in the model, which includes Ds, DPs, and MDs. Instead, we find that *N*(*t*)increases faster as *μ* increases leading to the occurrence of cancer more rapidly - a result that accords well with experimental observations^3^. In our model, the enhanced probability and rapid acquisition of cancer take place because the effect of DPs are compensated by the mini-drivers and drivers. In accord with expectations, we find that cancer progression rate is very slow as *μ* deceases as shown by the orange lines in Fig. 3. Therefore, cancer can be delayed if the mutation rate is decreased endogenously or exogenously. However, evolution would also be arrested or drastically slowed without mutation. From this perspective, cancer - a lethal disease, is a wicked by-product of evolution.

**Effect of death rate on population growth.** The growth in *N*(*t*)shown in Figs. 2–3 was obtained by assuming that the cell death rate increases linearly with *N* (Eq. (2)). The populations increase as a new driver mutation starts to get fixation resulting in an increase in the cell fitness by *s_d_*. Simultaneously, *D*(*N*) also increases as N becomes larger. In this model, *N* can only increase by a finite amount as each driver mutation gets fixation successfully. Other functions have also been considered for cell death rate in cancer research. In the following, we discuss two different functions for cell death to illustrate the influence of the choice of *D*(*N*) in tumor growth. A constant death rate is frequently utilized^33,35^, and we will consider this simplest function first. The evolution dynamics of cancers with Ds, MDs as well as DPs is illustrated in Fig. 4A. Here, *D*(*N*) is a constant with the assumption that it equals the birth rate at *t* = 0. Initially, we observe homeostasis, especially when *N*_0_ = 1000. This is followed by unbounded growth as the cells gain the first driver mutation irrespective of *N*_0_. It indicates that one driver mutation can lead to cancer, which in this instance occurs because the birth rate of cells is always much higher than the death rate after the first driver mutation gets fixation. The tumor grows to a macroscopic size in a very short time (Fig. 4A).

**FIG. 4.**
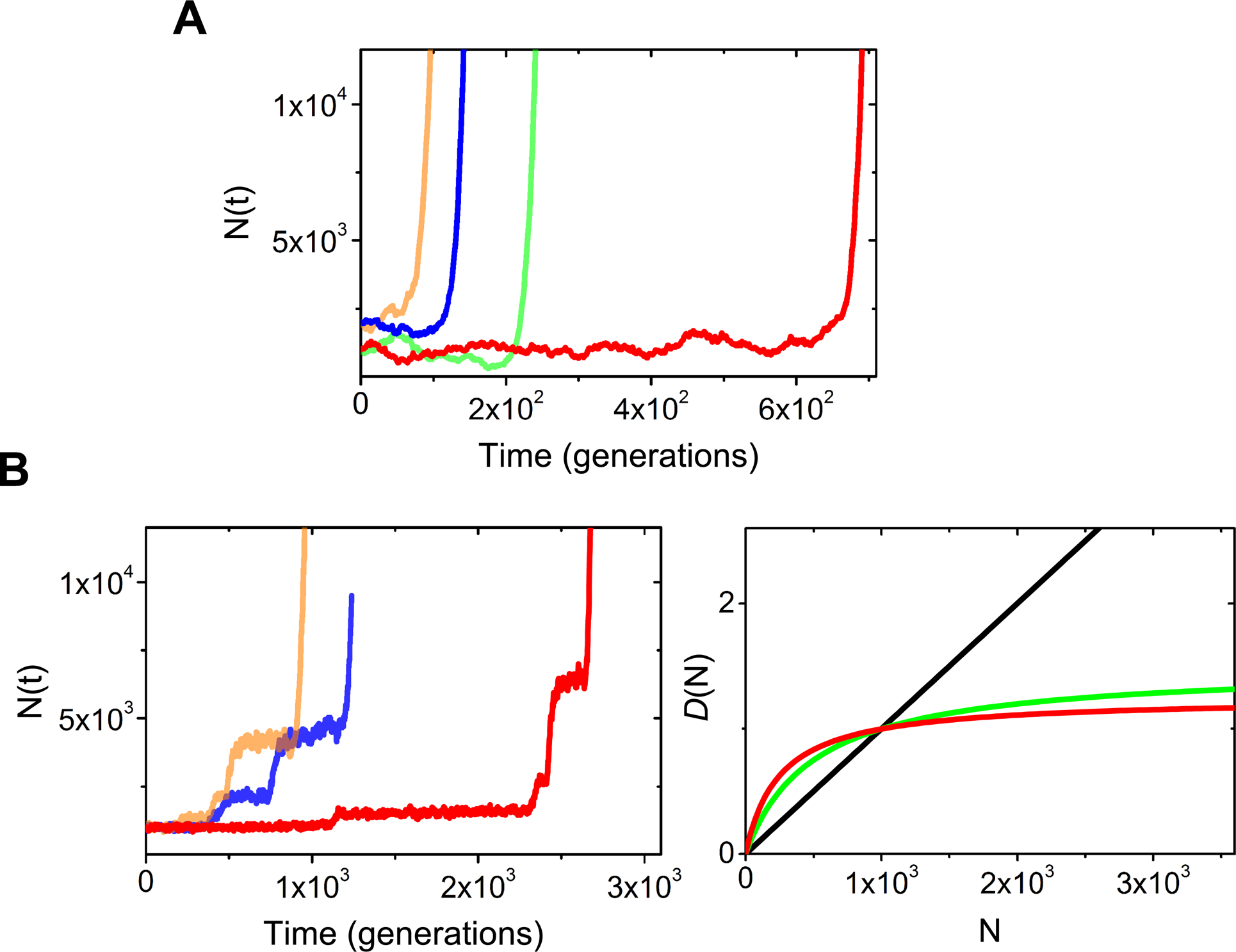
Dependence of growth dynamics on the functional form of death rate for a model with Ds, MDs, and DPs. (A) *D*(*N*) is a constant, and Initial sizes *N*_0_ = 1000 or 2000. (B) *D*(*N*) given by Eq. (4) with 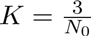 with *N*_0_· with *N*_0_ = 1000. The panel on the right shows a plot of *D*(*N*) (Eq. (4)) with 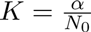 with *a* = 1(black), *α* = 3 (green), and *α* = 5 (red).

A constant value for cell death rate is not very realistic because cells are subject to spatial, resources and other constraints as the population grows to macroscopic sizes. Stimulated by a previous study^33^, we propose another function for *D*(*N*),

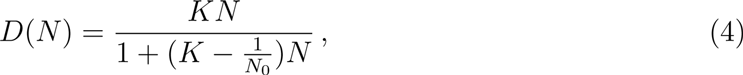

where *K* is a constant. If *K* = 1/*N*_0_, we recover Eq. (2). The death rate is still an increasing function of the system size *N*, but it increases slowly as *N* reaches large values. This is physically reasonable because certain driver mutations can promote the survival of cells by decreasing the death rate as discussed above.

An interesting phenomenon emerges when Eq. (4) is utilized for *D*(*N*). Initially, the growth dynamics (Fig. 4B) is similar as in Figs. 2–3, with homeostasis interrupted by periods of rapid population growth. However, the tumor transits from a bounded to an unbounded growth after several drivers (four in this example) accumulate. Therefore, only a small number of drivers is required for the occurrence of cancers^1,2,28,33^. The precise number of drivers is controlled by variables such as *K*, which might explain the varied risks for the occurrence of different cancers. It takes much longer time to develop cancer if a larger number of drivers are required for cancer progression (Fig. 4). Recently, it was found^36^ that African elephant which has a lower cancer mortality compared with human has at least 20 copies of TP53 (a tumor suppressor gene) while only one copy is present in humans. This observation provides a clue to resolve the Peto's paradox that large, long lived animals do not experience an increased risk of cancers^37^. Our simulations provide a qualitative explanation of Peto’s paradox based on one choice of N-dependent *D*(*N*).

### Simulations explain the number of drivers and passengers in cancer data

The number of drivers and passengers that accumulate in tumor cells under a variety of conditions is explored in Fig. S3-S6. The identification of driver mutations is a central topic in cancer research^3,38^ whereas the effects of passengers and mini-drivers have been ignored until recently^15,19,23^. The number of passengers increases as drivers accumulate in the same type of tumors^15,19^. In order to assess if the relation between driver and passenger predicted here is consistent with experiments, we analyzed data from six clinical sets (Fig. 5)^15,19,39,40^. In Fig. 5A, the filled star is for Glioblastomas (GBM) and the open star corresponds to data for pancreatic cancer. Four other types of cancer including melanoma, breast, colorectal, lung cancer are shown by open stars with different colors in Fig. 5B. A positive correlation is observed between the number of passengers and drivers. The clinical data shows that passengers accumulate faster than drivers although the precise number depends on the cancer type.

**FIG. 5.**
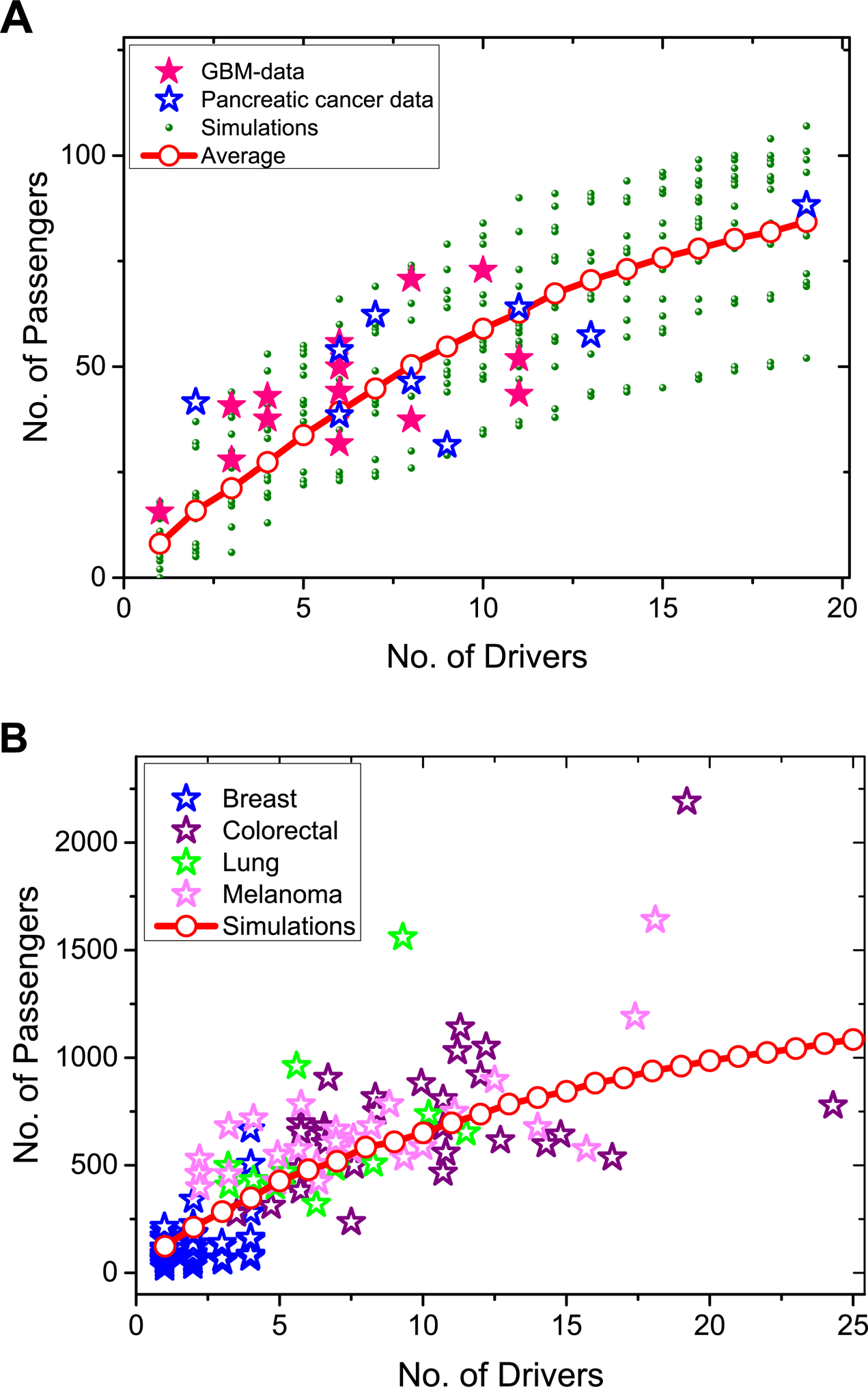
Comparison between clinical data and simulation results. The total number of passengers (DPs and MDs) versus the total number of drivers. (A) Two groups of clinical mutation data^15^ are shown in filled (GBM) and open stars (pancreatic cancer), respectively. Open circles are obtained by averaging over 40 trajectories. We plot 10 of them in green dots. The number *L*_*p*_ of passenger loci is 10^6^, and the values of other parameters are the same as in Fig. 2(C). (B) Four groups of clinical data^19^ are shown in open stars with different colors. Results from simulations are in open red circles. The parameters are the same as used in Fig. 2(C).

In comparing the simulation results and clinical data in Fig. 5 we assume that both DPs and MDs are regarded as passengers because of their relatively weak influence on tumor progression. The total number of passengers is the sum of DPs and MDs. The simulation results for the model including Ds, MDs, and DPs, illustrated by open circles in Fig. 5 describe all the six clinical mutation data well. The accumulation rate of passengers, as a function of the number of drivers, starts to decrease as the number of drivers reaches large values, which is consistent with previous studies^15^. It should be emphasized that passenger mutations were assumed to be neutral^15^, which is not the case in our study. Interestingly, there is a great deal of dispersion in the plot Fig. 5B. The fluctuations around the mean (red circles) is substantial. To the extent each curve in Fig. 5A (shown as green dots) represents the fate of a single cell population, the results show that there ought to be considerable heterogeneity in the plot of number of passengers versus drivers, as found in Fig. 5B.

### Deleterious driver results in population extinction at high mutation rates

So far, we have neglected the role of deleterious drivers (DDs) in the evolutionary growth model. Such mutations, which strongly reduce the fitness of tumor cells, could occur with some probability. Neglect of DDs is reasonable as long as the mutation rate (*μ*) is small. However, *μ* can be enhanced by specific (epi)genetic mutations, chromosomal instability (CIN), or carcinogens^3^. We show that DDs can play an important role as *μ* reaches high values.

We generalized the model to investigate the effect of DDs. We take the fitness disadvantage *s*_*dd*_ = *s_d_* = 0.1 for DDs. The number *L*_*dd*_ of DDs loci, which are associated with essential genes, is taken to be 10^4^ (see SI for an explanation of this choice). The evolution of *N*(*t*)at different *μ* values, plotted in the upper panel of Fig. 6, shows that the DDs do not influence the evolution of *N*(*t*)as long as *μ* ≪ 10^−4^. In this limit, the growth of *N*(*t*)is similar to that reported in Fig. 2C. Occurrence of cancer is more rapid as *μ* increases (see also Fig. 3B). The average number of DDs in a single tumor cell is negligible if *μ* ≪ 10^−4^ (lower panel in Fig. 6). Therefore, DDs can be neglected as long as *μ* is relatively small^19^. However, there is a dramatic change in the growth dynamics with tumor driven to extinction if *μ* > 10^−5^ as shown by the lines in orange and red in Fig. 6. This finding is not observed in a model including only Ds, MDs, and DPs (Fig. 2C). At such high mutation rates, the number of accumulated DDs in tumor cells increases rapidly, as illustrated by the open symbols in orange and red in Fig. 6. This results in a strong reduction of the fitness of tumor cells leading to the extinction of the population. The critical mutation rate constant *μ_c_* (> 10^−5^) decreases as *L*_*dd*_ increases (a high mutation rate for DDs can still be maintained).

**FIG. 6.**
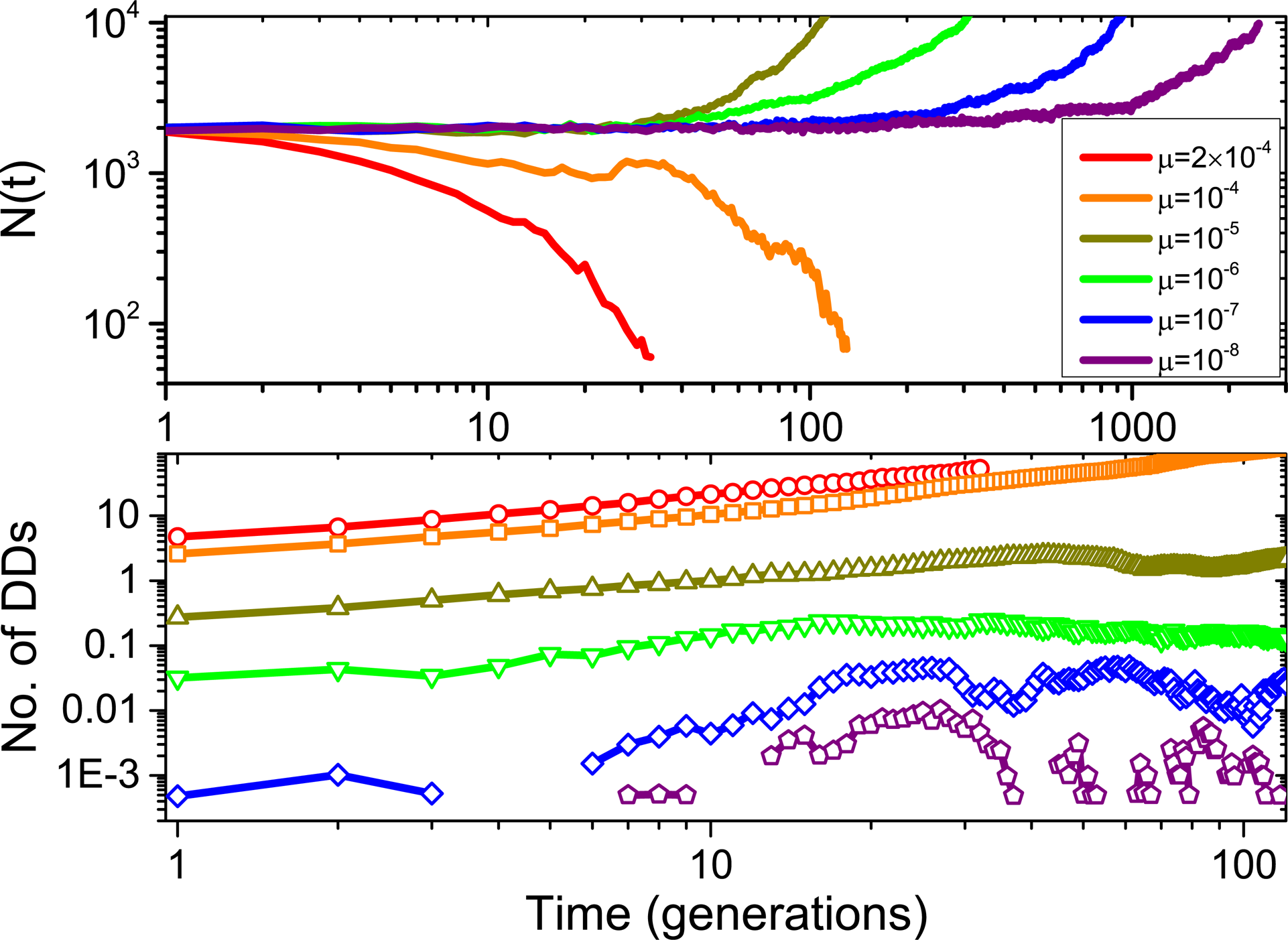
The dynamics of tumor evolution for a model with Ds, DDs, MDs, and DPs starting from initial size, *N*_0_ = 2000 at different *μ* values. The upper panel shows *N*(*t*)as a function of time. The values for the *μ* are in the inset. The lower panel shows the evolution of the average number of accumulated DDs per cell using *L_dd_* = 10^4^, *s*_*dd*_ = 0.1. The values of other parameters are from Table 1. Same color code is used for *μ* in the upper and lower panels.

The surprising finding that the extinction of *N*(*t*)at high *μ* due to the presence of DDs explains the counterintuitive correlation between high chromosomal instability (CIN) and improved prognosis for clinical outcome in a number of cancers^41,42^. Based on a score accounting for DNA based chromosomal complexity and CIN, which correlates with gene expression in ~ 2,000 breast tumors, it was found that the tumors with high CIN had greater prognosis compared with intermediate CIN. Our result, showing that only at high *μ* the presence of DDs is poorly tolerated by cancer cells, is consistent with the observed paradoxical relationships between high CIN and improved clinical outcome. These CIN scores can be calculated using microarray expression data available publicly, and hence it stands to reason that the genes for DDs can be identified. If so, enhancing their mutation rate might be a viable therapy for some forms of cancer.

Interestingly, it was also previously shown that the probability of cancer decreases if *μ* exceeds ~ 4 · 10^−9^ using a model with only drivers and deleterious passengers^19^. This observation also explains the paradox between improved cancer prognosis and high CIN. The decrease in cancer incidence in our model occurs at mutation rates > 10^−5^ provided DDs are taken into account. The difference of (3-4) orders of magnitude between our predictions and the one estimated elsewhere^19^ for the critical mutation rate will require experiments. The spontaneous random mutation rate at the single nucleotide level in normal cells^43^ is around 10^−9^ to 10^−10^ while the rate is nearly (2-4) orders of magnitude higher in tumor cells^44,45^ which is in accord with the values obtained in this work. We conjecture (further experiments are needed) that *μ* needs to exceed 10^−5^ for a decrease in the probability of cancer. It should be emphasized that it is likely that high mutation rates trigger T-cell response that results in control of cancer growth rather than decreasing the fitness of cancer cells. Such a mechanism could also explain the successes of immunotherapy^46^ in the treatment of high mutation rate cancers such as melanoma. These two mechanisms are not mutually exclusive and could operate in concert.

## DISCUSSION

We investigated the interplay between rare driver mutations that confer distinct fitness advantage to cancer cells, and the numerous but weak mini-drivers and deleterious passengers in the evolution of cancer. These conflicting effects, such as the mini tug-of-war between the MDs and DPs, result in rich dynamics in tumor growth. Simulations based on the new evolutionary model show that both DPs and MDs profoundly impact the growth dynamics of tumors. By including MDs, we find that the dynamics of population growth exhibit intermittency with periods of homeostasis (plateaus in *N*(*t*)) followed by bursts in the growth. If DPs are totally ignored and dynamics only with Ds and MDs are considered the population grows continuously with varying growth rates (see Fig. S8 in the SI). By including Ds, MDs, and DPs we find population homeostasis, which accords well with the observation that the rapid growth of tumors is often interrupted by periods of dormancy^33,34^. The effects of deleterious drivers are negligible as long as the mutation rate does not reach very high values. However, DDs accumulate in tumor cells rapidly as the mutation rate becomes high (> 10^−5^). When this occurs, the tumor size decreases because of the strong deleterious effect of DDs. This explains the paradoxical phenomenon that high CIN correlates with improved prognosis in several types of cancer. The optimal mutation rate (*μ_c_* ~ 4 · 10^−9^) above which probability of cancer occurrence is low, predicted by the tug-of-war model, is significantly lower than our predictions (*μ_c_* > 10^−5^).

The new model, which includes mini-drivers and deleterious drivers captures many dynamical features of cancers successfully. The model can also be used to investigate other properties of cancers such as the fixation process of mutations, and their distributions in cancer cells. However, there are limitations to almost all of the evolutionary models. It is well known that cancer cells often show resistance to chemotherapy and radiation therapy, which strongly reduces the effect of treatments resulting to relapse. It is likely that cancer heterogeneity^47^ could be the reason for the lack of efficacy of standard treatments. Cancer cells are usually composed of many subpopulations with distinct genetic and phenotypic variations. Therefore, the assumption of well-mixed population has to be relaxed to account for heterogeneity, a hallmark of many cancers, whose importance is a continued focus of cancer research. This requires inclusion of interactions between subclones^48,49^, and spatial structures of tumors^35,50^.

## Acknowledgments

We are indebted to Shaon Chakraborty for several pertinent comments and suggestions. We are grateful to Abdul Nasser, Mauro Mugnai, Himadri Samanta, Sumit Sinha, Huong Vu for discussions and comments on the manuscript. This work, initiated when the authors were at the Institute for Physical Science and Technology at the University of Maryland, is supported by the National Science Foundation (CHE 16-36424).

